# Utilization of Eco-friendly Imidazolium-Based Ionic Liquids for Gluten Extraction: Investigating the Influence of Side Chains and Anions

**DOI:** 10.1101/2023.09.17.558121

**Authors:** Wen-Hao Chen, Yu-Cheng Hsiao

## Abstract

Gluten is a well-known food allergen globally, capable of triggering immune responses in both celiac and non-celiac gluten-sensitive individuals. Gluten comprises two major proteins: glutenin and gliadin. Gliadin, in particular, possesses a unique hydrophobic amino acid sequence. The Food and Drug Administration asserts that the toxicity of gliadin cannot be eliminated through fermentation and hydrolysis processes. A common approach to addressing gluten allergies is to adopt a gluten-free diet. However, the hydrophobic nature of gluten makes its detection challenging. Analysts often resort to using organic solvents or employing multiple procedures to denature gluten for extraction. It’s worth noting that while organic solvents can rapidly extract gluten from a sample, they may also alter antibodies, leading to erroneous bio-test results.

Ionic liquid (IL) is a highly adaptable green chemical compound consisting of organic salts. We modified imidazolium, a cationic structure, with various carbon side chain lengths (C=0, 1, 3, 5, 7, 9, and 12), and combined it with organic and inorganic anions (e.g., OMs-, Cl-, F-, NO¬3-, HSO4-, and H2PO4-). We used different IL-to-water ratios to assess gluten solubility. We measured the solubility of gliadin in various imidazolium ILs and conducted kinetic studies on the dissolution of gliadin in 1% [C5DMIM][OMs]aq. Additionally, circular dichroism (CD) spectroscopy and enzyme-linked immunosorbent assay (ELISA) were employed to evaluate the structural changes in gliadin and its interaction with antibodies after treatment with 1% [C5DMIM][OMs]aq. An XTT assay was conducted to assess the cytotoxicity of [C5DMIM][OMs]aq on N2a cells.

Our research findings indicate that 1% [C5DMIM][OMs]aq demonstrated excellent gluten solubility, dissolving more than 3000 ppm of gluten within 5 minutes. Importantly, [C5DMIM][OMs]aq did not disrupt the gluten structure, did not impede antibody binding to gluten, and exhibited no cell toxicity. This report highlights [C5DMIM][OMs] as a promising extraction solution for gluten detection.

## Introduction

Around 0.5%∼6% of people worldwide have celiac disease or non-celiac gluten sensitivity (NCGS)^1–5^. Celiac disease and NCGS are food allergy diseases, and the reason comes from gluten-induced immune responses that attack the body^6, 7^. Gluten allergy symptoms include bloating, chronic diarrhea, depression, anxiety, etc. It raises the risks of enteropathy-associated T-cell lymphoma (EATL), non-Hodgkin’s lymphoma, and adenocarcinoma of the small intestines when celiac patients do not control their gluten intake^8–10^.

Gluten is a major food allergen worldwide, and it is comprised of two major proteins: glutenin and gliadin^11^. Glutenin is water soluble, while gliadin is insoluble in water. Some research claimed that gliadin appears to be the primary cause of celiac disease. Gliadin contains gliadin groups (alpha/beta, omega, and gamma) and gliadin subunits (high-(HMW) and low-molecular weight (LMW)). Regardless of the kind of gliadin, gliadins are capable of aggregating into larger oligomers and interacting with other gluten proteins due to large hydrophobic sections^12, 13^, poly-Q, and repetitive sequences^14–16^. These sections are likely to aggregate hydrophobically^17^, separate in the liquid-liquid phase, potentially form β-sheet aggregates^18^, or simply become entangled by their structural properties. Detection of gluten is difficult due to the effects of hydrophobicity and aggregation. Analysts need to use organic solvents or spend more time pretreating a sample to extract gliadin from a sample. Recently, scientists have tried to use certain enzymes to hydrolyze gluten to decrease its toxicity^19–21^, but still no useful process has been found to decrease the toxicity ^22, 23^ according to Food and Drug Administration (FDA) publication in 2022.

Although food technology keeps progressing, the only good way to so far treat a gluten allergy is to avoid consuming gluten. Generally for gluten analysis, an enzyme-linked immunosorbent assay (ELISA) is the main gluten detection system. An ELISA has high specificity for gluten antibodies. However, gluten is difficult to dissolve in water solutions, especially in phosphate-buffered saline (PBS) ^24^. Analysts need to pretreat samples with 75% alcohol or spend time heating the sample in solutions ^25–27^. Alcohol is detrimental to antibodies, as it can induce antibody mutations causing them to lose their function. Therefore, the analyst needs to transfer alcohol to buffer system (e.g. PBS buffer) to reduce its effects on the antibody. It is a complicated process which takes time to pretreat samples, regardless of the process chosen.

Ionic liquids (ILs) are synthesized by organic cations and organic/inorganic anions^28, 29^. In most cases, the cation or anion is tetraalkyl ammonium, tetraalkyl phosphonium, imidazolium, cholinium, or pyridinium. ILs function can be varied by modifying side chains, and they can have different physical and chemical effects by changing the anions^29^. ILs can be applied to heavy metals^30^, little molecular^31^ extraction and also to stable protein structures^32^. Based on the ability to modify ILs, we designed and synthesized different lengths of side chains and anions for rapid extraction of gluten from samples. In this report, we used imidazolium as the major structure, modified with adding different lengths of carbon side chains for interactions with hydrophobic sides of gluten, and changed the anion to increase the solubility of gluten.

ILs we synthesized provide the interaction to gliadin protein. Ionic forces come for the salt of hydration radius interact with a protein. Van der Waals forces come for interactions with carbon lengths. Generally, the van der Waals force is increased with length. Since gliadin has special amino acid sequences like poly glutamine and continuous poly proline, these special amino acid sequences induce protein-water insolubility and cause the fiber structure to easily aggregate^33, 34^. For this special amino acid sequence, we synthesize a suitable side chain of IL to bind to the gliadin protein, to make gliadin soluble in water by for a safe and effective analysis to galidin detection.

## Results and Discussion

### Solubility of gluten in [C5DMIM][OMs]_aq_

Ionic solutions of [pentyl dimethyl imidazolium][methyl sulfonic] ([C5DMIM][OMs]) at concentrations of 0.05, 0.1, 1, 2, 5, and 10 wt% in water were prepared in sample vials. Three grams of gliadin (an excess amount) was added to 10 mL of each [C5DMIM][OMs] ionic solution, mixed homogenously by stirring for 30 min, and allowed to sit for 5 min for the test example. All sample solutions were centrifuged at 8500 rpm for 3 min and filtered through a 0.22-μm-pore size filter.

Water (0 wt%) was used as the control group. After that, a Wheat/Gluten (Gliadin) ELISA Kit (Crystal Chem, AOAC no. 011804) was used to determine the concentration and solubility of gliadin. Since the [examination/limit?] of the Wheat/Gluten (Gliadin) ELISA Kit was about 100 ppm, the gliadin concentration was determined by diluting sample solutions by at least 50-fold with a corresponding ionic solution in order to calculate their gliadin concentrations. As shown in Fig. 1, gliadin solubility increased when the percentage of [C5DMIM][OMs] was raised from 0% to 0.1%, and the maximum value of gliadin solubility was at 3000 ppm of 1% [C5DMIM][OMs]_aq_. There was no significant difference in gliadin solubility when [C5DMIM][OMs]_aq_ was raised to 10%.

**Figure 1.**
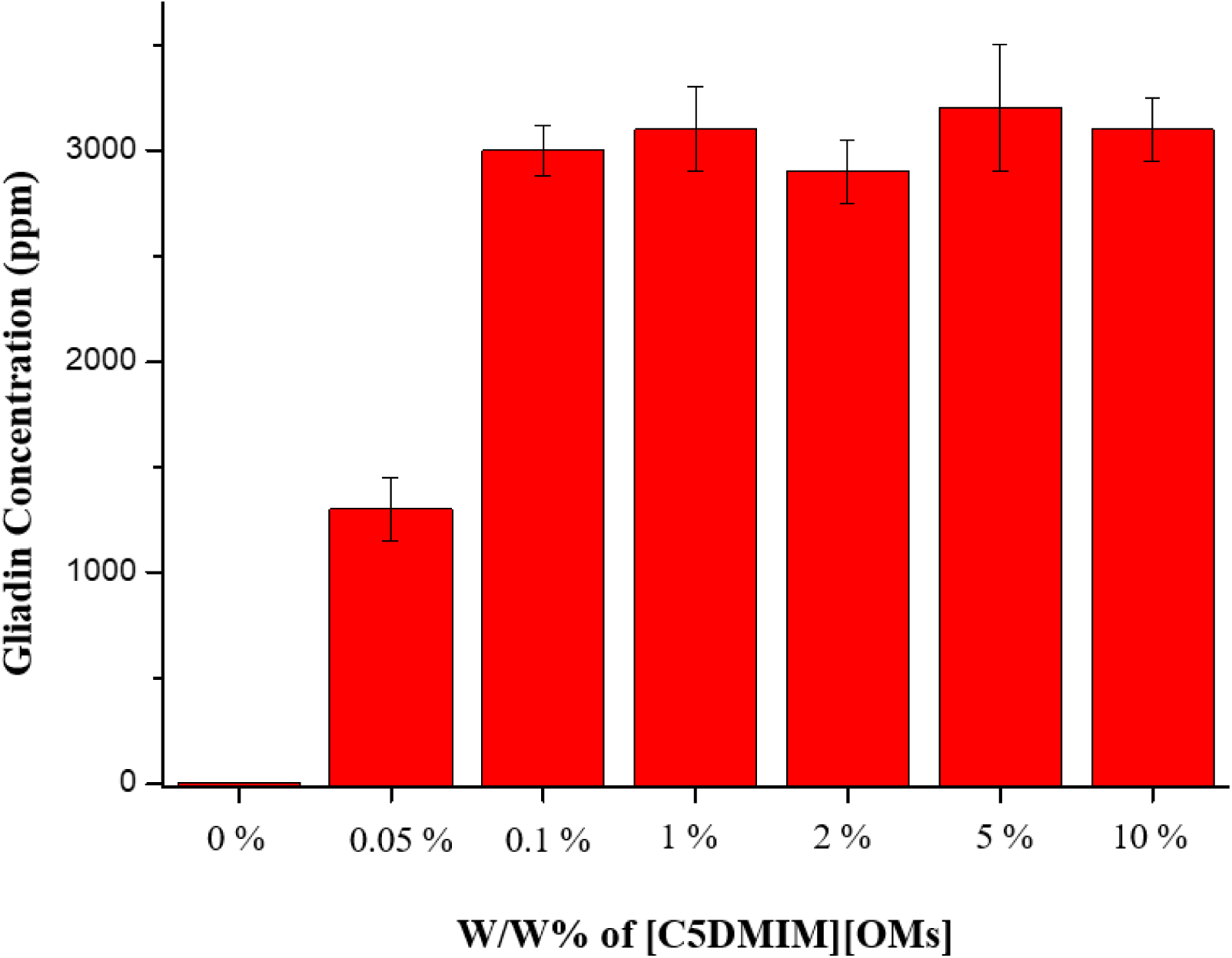
Gliadin solubility in the [C5DMIM][OMs] water solution.

### Side chain effects on gluten solubility in 1 wt% of an IL water solution

The special function originates from modifications of the IL side chains. In this report, we synthesized different side chain lengths to test the IL solubility of gluten. In recent years, some reports showed that longer side chains and anions of ILs could penetrate cell membranes and damage cells and macrobiotics^35, 36^. For this reason, we tried to synthesize a more environmentally friendly ILs and applied them to gluten extraction.

We synthesized and prepared 1% IL water solutions of {[hydrogen dimethyl imidazolium][methyl sulfonic] ([HDMIM][OMs]), [trimethyl imidazolium][methyl sulfonic] ([TMIM][OMs]), [propyl dimethyl imidazolium][methyl sulfonic] ([C3DMIM][OMs]), [C5DMIM][OMs],[heptenyl dimethyl imidazolium][methyl sulfonic] ([C7DMIM][OMs]), [nonathyl dimethyl imidazolium][methyl sulfonic] ([C9DMIM[OMs]], and [dodecathyl dimethyl imidazolium][methyl sulfonic] ([C12DMIM][OMs])) in sample vials. Three grams of gluten was added to 10 mL of 1 wt% of each IL solution, homogenously mixed by stirring for 30 min, allowed to sit for 5 min, used a 0.22-µm filter to remove precipitates, and measured the gliadin concentration with a gliadin ELISA kit.

As shown in Fig. 2, gliadin had a higher solubility in the [C5DMPIM][OMs], [C7DMIM][OMs], [C9DMIM][OMs], and [C12DMIM][OMs] IL solutions, in which the corresponding groups attached to the 3-position of 1,2-dimethyl imidazolium had different carbon numbers of 5, 7, 9, and 12, respectively. The [C9DMIM][OMS] and [C12DMIM][OMs] ILs had lower solubilities but higher viscosities, which might be disadvantageous. However, decreasing the length of the carbon chains also reduced toxicity toward the environment^35, 36^.

**Figure 2.**
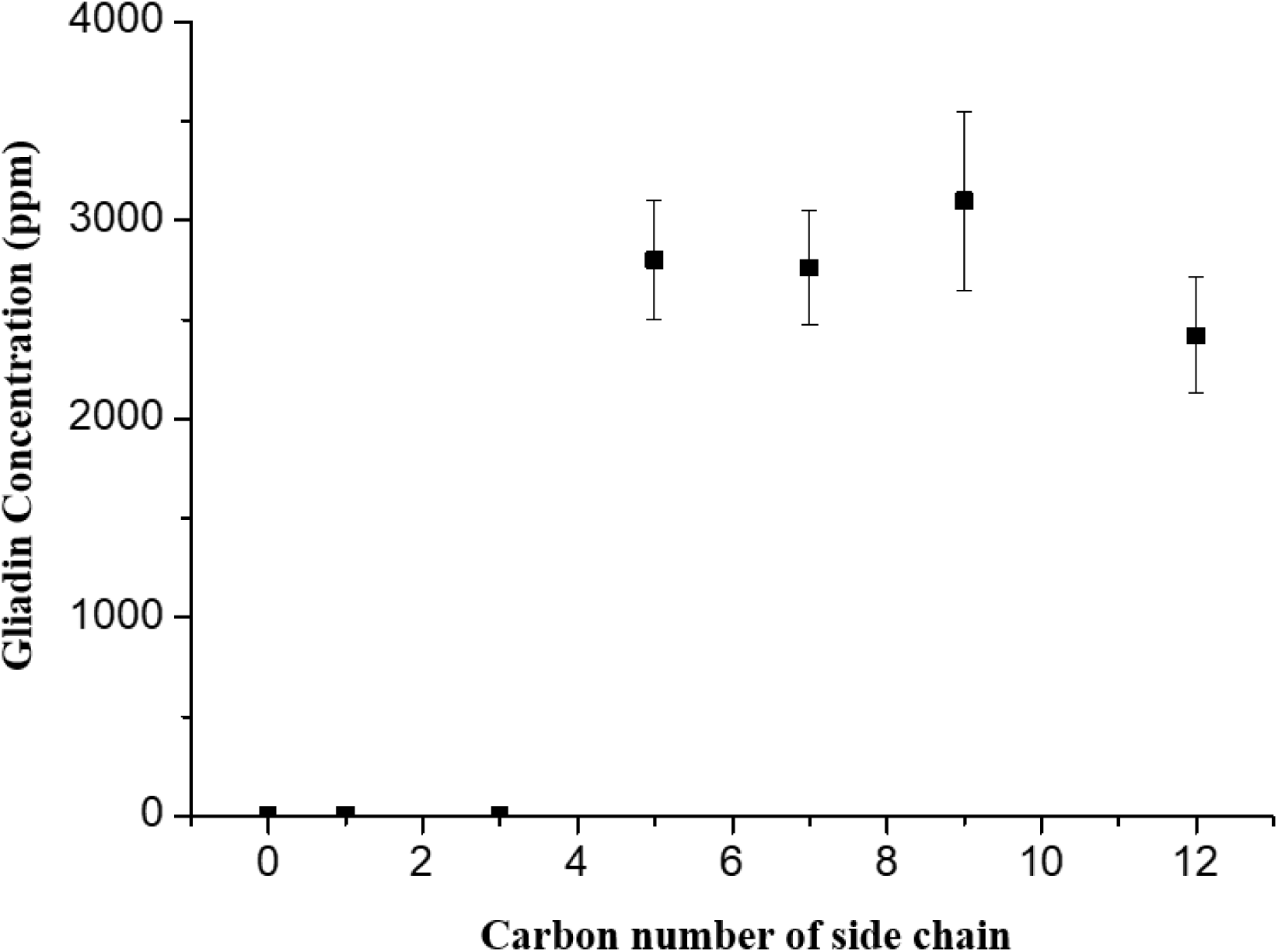
Side chain effect of imidazolium with methane sulfonic acid anions on the gliadin solubility test.

### Solubility of gliadin in 1 wt% ionic solutions of ionic compounds with different anions

Different anions will have different effects in ILs. The hydrophobicity, hydrophilicity, and melting and boiling points can be controlled by different anions of ILs^29, 37^. In this report, we changed the anions to test the gluten solubility of ILs. We synthesized and prepared 1% IL water solutions of [C5DMIM][Cl], [C5DMIM][F], [C5DMIM][OMs], [C5DMIM][HSO_4_], [C5DMIM][NO_3_], and [C5DMIM][H_2_PO_4_] in sample vials, and 3 g of gluten was added to 10 mL of a 1 wt% IL solution, mixed homogenously by stirring for 30 min, allowed to sit for 5 min, used a 0.22-µm filter to remove precipitates, and measured the gliadin concentration with a gliadin ELISA kit. As shown in Fig. 3, the solubility of gliadin in the solutions of ionic compounds with halogen anions was low, and the ionic compound with [OMs^-^] anions produced the greatest gliadin solubility. Gliadin solubility still had a good effect with certain inorganic anions (HSO_4_^-^, NO_3_^-^, and H_2_PO_4_^-^).

**Figure 3.**
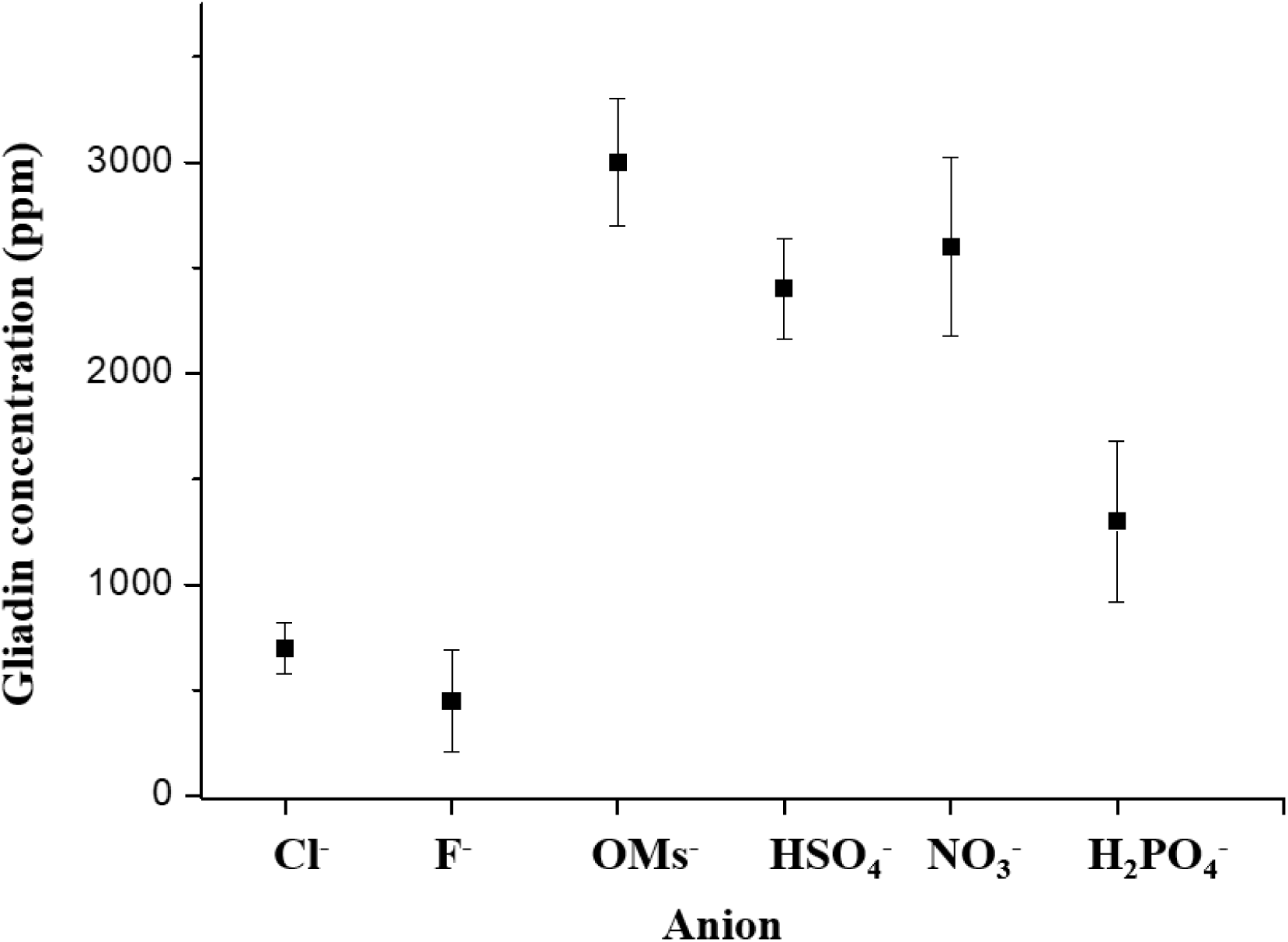
Anion effects on the gliadin solubility test with [C5DMIM].

### Kinetics of the solubility of gliadin in 1 wt% [C5DMIM][OMs]_aq_

We prepared 1% [C5DMIM][OMs]_aq_ and PBS in sample vials, separately added 3 g of gliadin into the vials, homogenously mixed it by shaking the vials, and allowed it to sit for various intervals (0.5, 1, 2, 3, 5, 10, 20, and 30 min). We used a 0.22-µm filter to remove precipitates, and the gliadin solubility was measured with a gliadin ELISA kit. As the control test, PBS was selected to substitute for [C5DMIM][OMs], and the same process as for pretreating the gluten sample was carried out. We used the ELISA to measure the gliadin concentration after pretreatment.

PBS is an a well-known buffer system for gliadin extraction^28^. As shown in Fig. 4, the solubility of gliadin with PBS extraction was <5 ppm with a longer time for extraction (red dot), and the solubility of gliadin was >700 ppm in 30 s after [C5DMIM][OMs]_aq_ extraction, and the solubility of gliadin reached its maximum in 5 min (black square).

**Figure 4.**
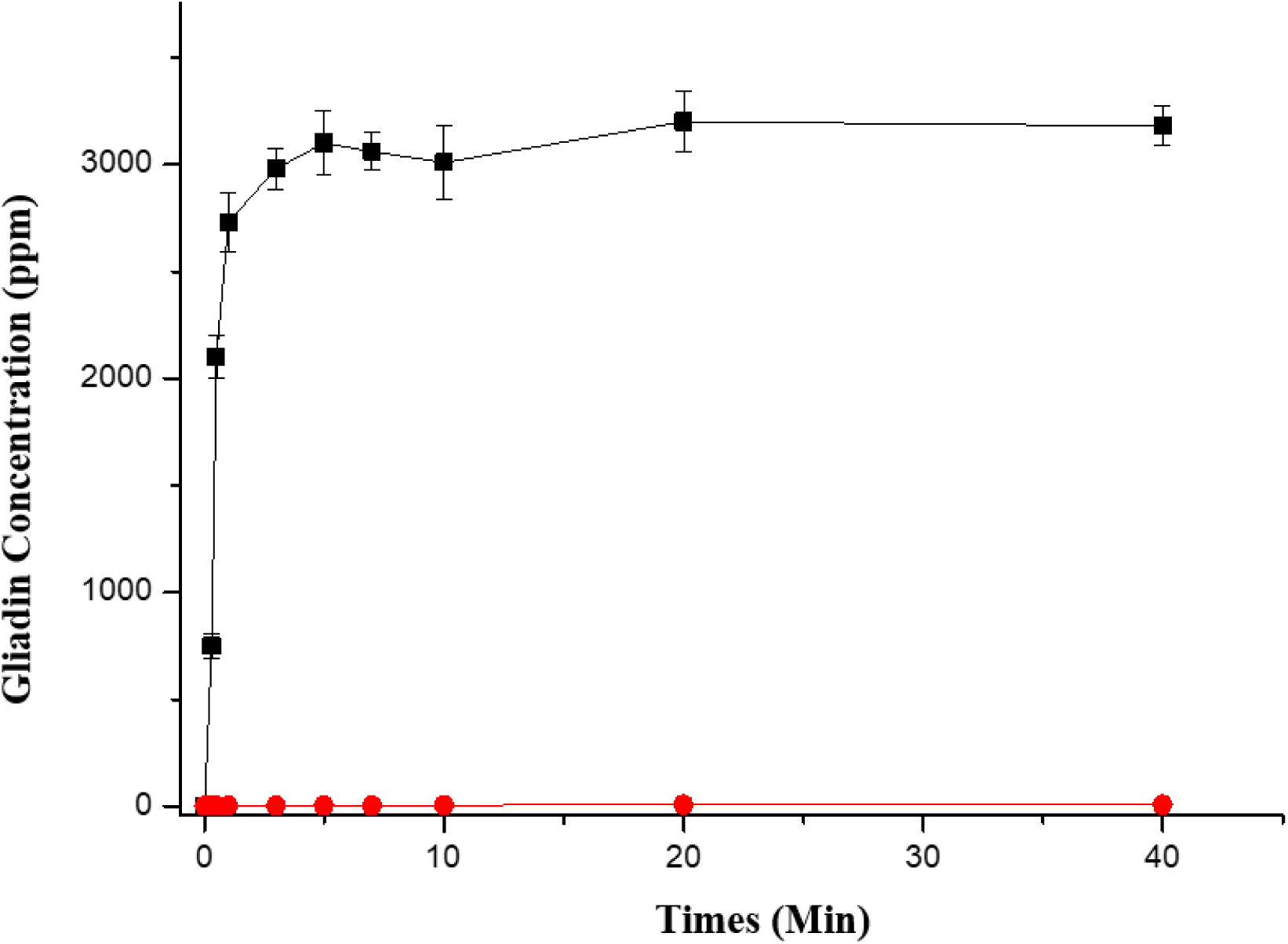
Kinetic curve of gliadin dissolved in 1% [C5DMIM][OMs]aq (black squares) and PBS (red dots).

### Structure of gluten with/without IL extraction

Circular dichroism (CD) spectroscopy is very sensitive to the secondary structure of polypeptides and proteins. It is usually used to study the secondary structure (alpha helix/beta sheet) of peptides or proteins. In this report, we used CD spectra to measure the secondary structure of gliadin with/without [C5DMIM][OMs] extraction. A gliadin sample without an IL solution (20 μM) was prepared from a standard gliadin solution (1 mg/ml) in PBS, which was directly prepared using commercially available gliadin (Leadgene). For the gliadin sample without an IL solution (20 μM), 1 ml of the solvent was removed by a vacuum, and then 1 ml of 1% of the IL ([C5DMIM][OMs]) solution was added to prepare the sample with IL extraction.

A CD Spectrometer (Jasco J-815) was used for the structural analysis of the above-mentioned sample solutions obtained by gliadin-rice noodle sample extraction and a commercially available gliadin product. As shown in Fig. 5, a beta sheet structure was evident between the extracted gliadin and commercially available gliadin (not extracted with the currently conceived ionic solution). Figure 5 shows that gliadin wt1% [C5DMIM][OMs] (red line) was less than gliadin with 1% [C5DMIM][OMs] (black line) at 220∼230 nm, which indicates that the IL interacted with gliadin and changed a little bit of its structure. But, generally, the CD spectra were similar between gliadin with and that without [C5DMIM][OMS] treatment, which means that the secondary structure of gliadin did not significantly differ after 1% [C5DMIM][OMs] treatment of gliadin.

**Figure 5.**
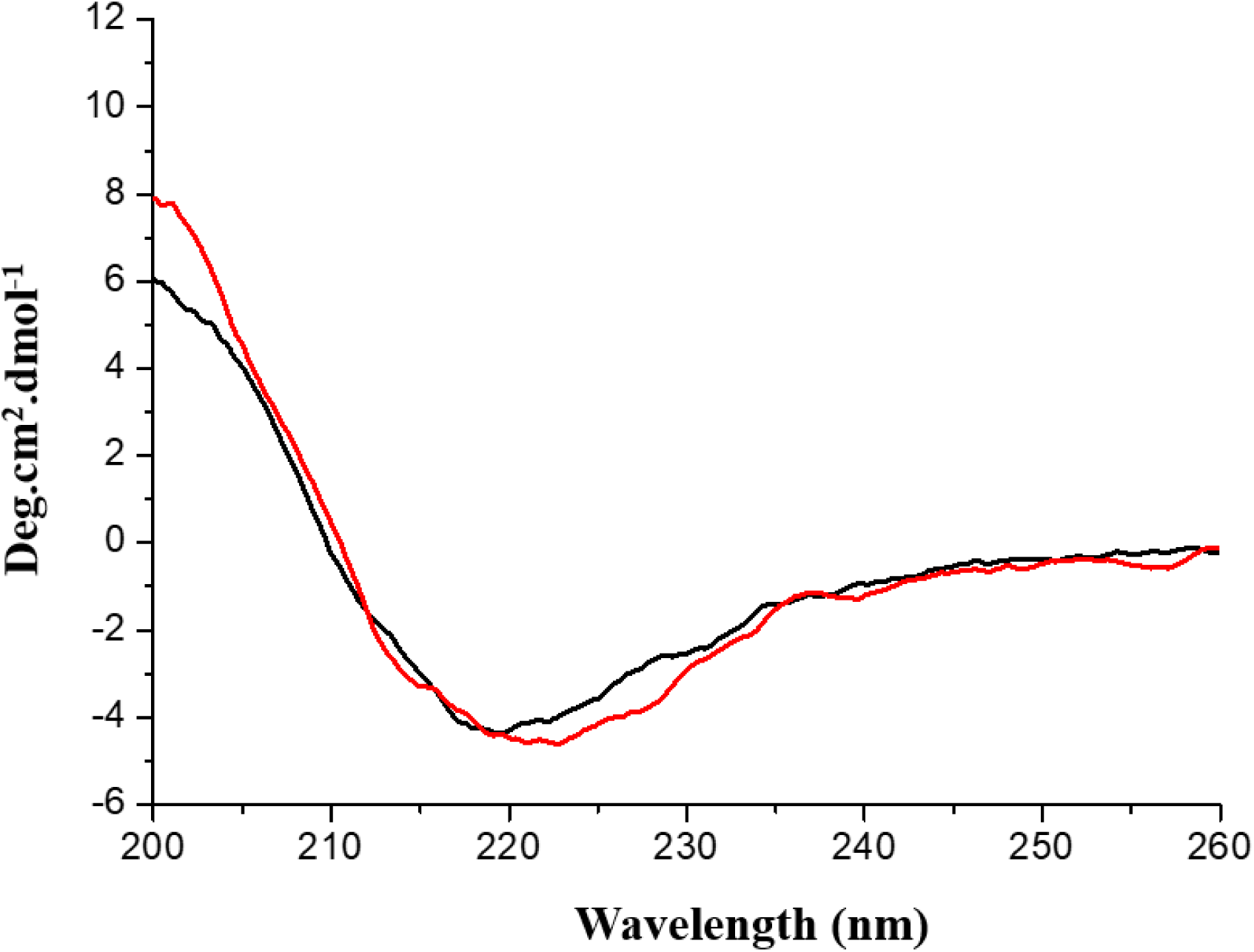
CD spectra of gliadin with (black line) and without (red line) [C5DMIM][OMs] treatment.

### Biochemical analysis of gliadin resolved in 1 wt% ionic solutions of 1 wt% [C5DMIM][OMs]

Biosensors are a very important technology globally, and this technology is widely used for disease diagnosis, detection of microorganisms and viruses, and all kinds of biomarker tests. The ELISA is widely used in biosensor issues, because it is specific and sensitive for target proteins according to the antibodies used. Certainly, the ELISA is the major biosensor for gluten now.

In this report, we used an ELISA to test gliadin binding with an antibody after 1% [C5DMIM][OMs] and control (1% sodium dodecylsulfate (SDS)) treatment. The 1% IL solution of [C5DMIM][OMs] in water was first prepared. Specific amounts of gliadin were added to the 1 wt% IL solution of [C5DMIM][OMs] to prepare gliadin sample solutions at 1, 2, 3, 5, 7.5, 10, 20, and 40 ppm of gliadin. After that, gliadin concentrations were confirmed by a Wheat/Gluten (Gliadin) ELISA kit.

A 96-well empty ELISA plate was coated with 1 μg/mL of anti-gliadin mouse immunoglobulin G (IgG; 2F, purchased from Antaimmu) and then blocked with 1% bovine serum albumin (BSA) to obtain an ELISA plate for gliadin examination. Gliadin sample solutions were respectively added to wells of the ELISA plate to react for 5 min, and then the wells were washed by PBS. After removing excess solution, 0.1 μg/mL anti-gliadin human IgA-horseradish peroxidase (HRP) (anti-gliadin human IgA 3B7 purchased from Leadgene, and HRP added by the MagicLink™ HRP Antibody Conjugation Kit) was added to react for 15 min, the plate was washed, TMB/H_2_O_2_ reagent (T3854, purchased from TCI) was added to react for 10 min, and a 0.5 mol/L sulfuric acid aqueous solution was added to terminate the reaction. The absorbance at 450 nm of all wells on the plate was examined by an ELISA reader (TECAN Infinite 200 PRO).

As shown in Fig. 5, the regression line had an *R*^2^ value of 0.9942. This indicates that the [C5DMIN][OMs]aq interacted with gliadin to raise the solubility, and the antibody could still bind with gliadin after [C5DMIM][OMs] interaction. For the control test (1% SDS for extraction), the absorbance was lower than the [C5DMIM][OMs] model, and its curve had a maximum at 20 ppm, which probably came from the strong cleaning effect of SDS to remove the antibody original coated onto the plate.

### Gluten recovery rate study with [C5DMIM][OMs]

The recovery rate is an important factor of extraction systems, and it will show the effect of extracting the target in a sample. In this report, we tested the recovery of gliadin by 1% [C5DMIM][OMs]. Rice noodles are a well-known food without gluten. In this report, we chose rice noodles as a substrate to test the recovery of gliadin by 1%[C5DMIM][OMs]. A 1 wt% IL solution of [C5DMIM][OMs] in water was first prepared. Hydrophobic proteins were traditionally extracted by an alcohol solution, so a 75 wt% ethanol solution was also prepared for this test.

Three grams of bread flour (Blue Jacket Strong Flour, Lien Hwa Milling) was mixed with 10 mL of 75 wt% ethanol for extraction at room temperature for 5 min to produce standard sample solutions. The sample solutions were centrifuged at 8500 rpm for 3 min and filtered with a 0.22-μm-pore size filter to obtain filtered samples. After filtration, the Wheat/Gluten (Gliadin) ELISA Kit (Crystal Chem, AOAC no. 011804) was used to determine gliadin concentrations in the filtered samples. The gliadin concentration was determined by diluting the sample solutions by at least 50-fold with the corresponding ionic solution, in order to calculate their gliadin concentrations. In both groups of 75 wt% ethanol extraction, gliadin was higher than 2000 ppm.

In addition, 40 g of dried gluten-free rice noodles (Organic Rice Noodles, Yuan Shun Food) was first soaked in water to rehydrate the noodles. After rehydration, the rice noodles were drained and soaked in 10 mL of a 200 ppm solution of gliadin in 100% ethanol in a container at room temperature to produce gliadin-rice noodles. The gluten-free rice noodles had a very large specific surface area (total surface area per unit of bulk volume), and most gliadin was adsorbed by the rice noodles.

After the ethanol solvent was evaporated, the gliadin-rice noodles were lyophilized in a container to give a gliadin-rice noodle sample. Ten milliliters of a 1 wt% IL solution of [C5DMIM][OMs] (equal volume with the 200 ppm gliadin solution in ethanol) was added to the lyophilized gliadin-rice noodles for extraction at room temperature for 5 min to produce test sample solutions. The sample solutions were centrifuged at 8500 rpm for 3 min and filtered with a 0.22-μm-pore size filter to obtain filtered samples. All groups were repeated five times. The concentration of gliadin was also determined with the Wheat/Gluten (Gliadin) ELISA Kit (Crystal Chem, AOAC no. 011804), wherein at least 5-fold diluted sample solutions with a 1 wt% IL solution of [C5DMIM][OMs] were used. The average recovery rate of the 1 wt% ionic solution of [C5DMIM][OMs] was 97.5%. The recovery rate was calculated using 200 ppm as 100%.

**Table 1.**
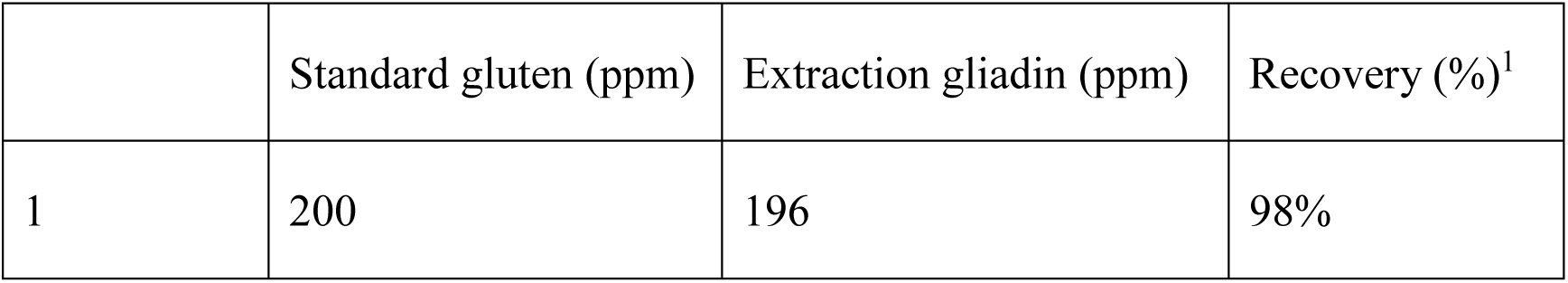
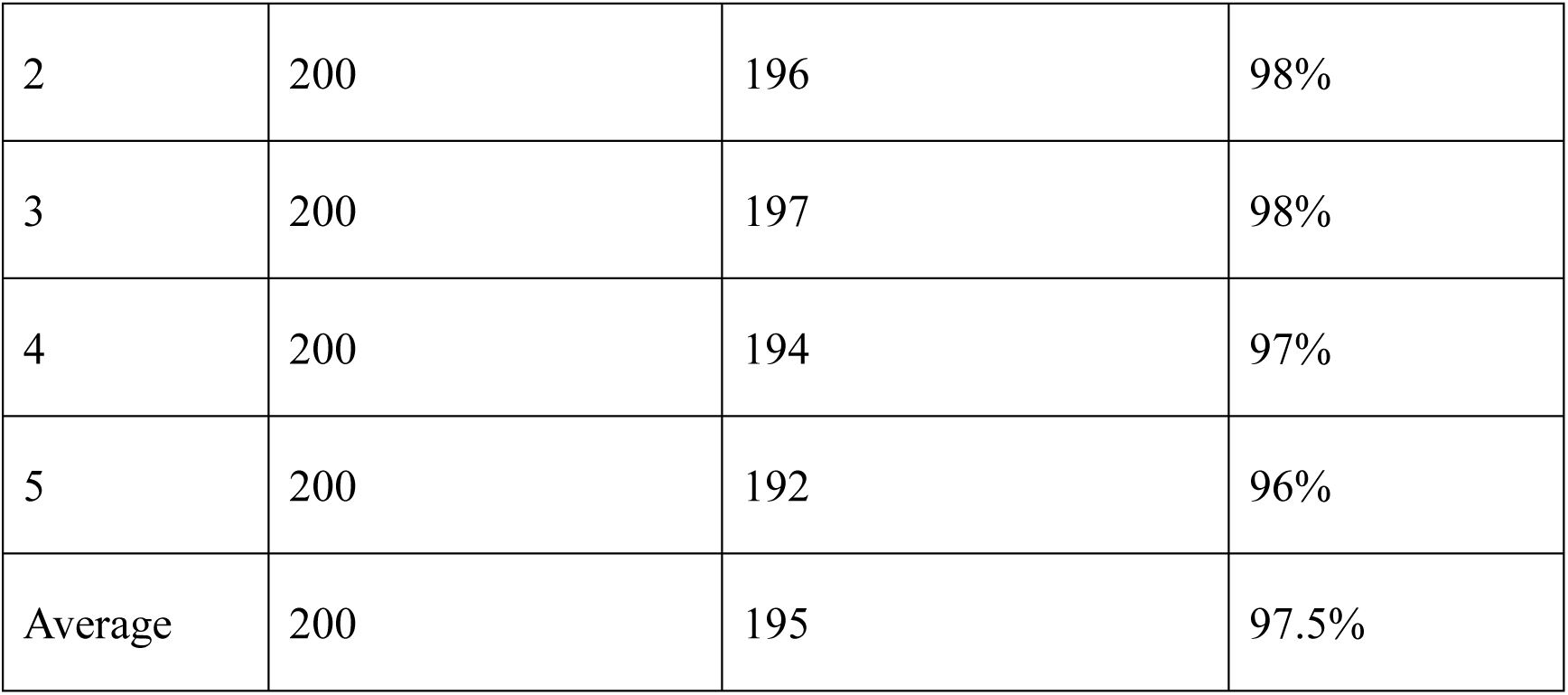
Percent Recovery = [extraction of gluten by the ionic liquid solution/standard gluten concentration] x 100%.

|  | Standard gluten (ppm) | Extraction gliadin (ppm) | Recovery (%)<sup>1</sup> |
| --- | --- | --- | --- |
| 1 | 200 | 196 | 98% |
| 2 | 200 | 196 | 98% |
| 3 | 200 | 197 | 98% |
| 4 | 200 | 194 | 97% |
| 5 | 200 | 192 | 96% |
| Average | 200 | 195 | 97.5% |

### Biocompatibility of IL solutions of [C5DMIM][OMs]

In recent years, the IL structure was reported to have cell toxicity towards cells and bacteria due to its long side chain effect (ref). For this reason, we tested the cell toxicity of [C5DMIM][OMs] in this report. XTT is commonly used to test nonradioactive quantification of cellular proliferation, viability, and cytotoxicity. The sample material was either adherent or suspended cells cultured in 96-well microplates. An increase in the number of living cells resulted in an increase in the overall activity of mitochondrial dehydrogenase in the sample. This increase was directly correlated with the amount of orange formazan formed, as monitored by the absorbance.

In this report, we chose mouse N2a neuroblastoma cells (N2a cells) to test the toxicity of [C5DMIM][OMS]. N2a cells were cultured and maintained in high-glucose Dulbecco’s modified Eagle medium (DMEM) with L-glutamine and sodium pyruvate (DMEM-HPA-P10, Capricorn Scientific) containing 10% fetal bovine serum (FBS). N2a cells were seeded in six-well plates at a density of 2×10^5^ cells/well overnight. Then the medium was replaced with test high-glucose DMEM medium containing 0.1, 1, 2, 5, or 10 wt% [C5DMIM][OMs], and cells were incubated in an incubator (37 °C, with a 5% CO_2_ humidified atmosphere) for 6 h. The test medium was removed, the cells were washed, and fresh high-glucose DMEM containing 10% FBS was added and incubated for another 12 h. After that, cells were subjected to an XTT test (X12223, purchased from Thermo Fisher) to determine the cell survival rate.

As shown in Fig. 7, survival rates of cells in the [C5DMIM][OMs] ionic solutions at all concentrations were high than 97%. This indicates that the ionic compounds in the present formulation had good biocompatibility.

**Figure 6.**
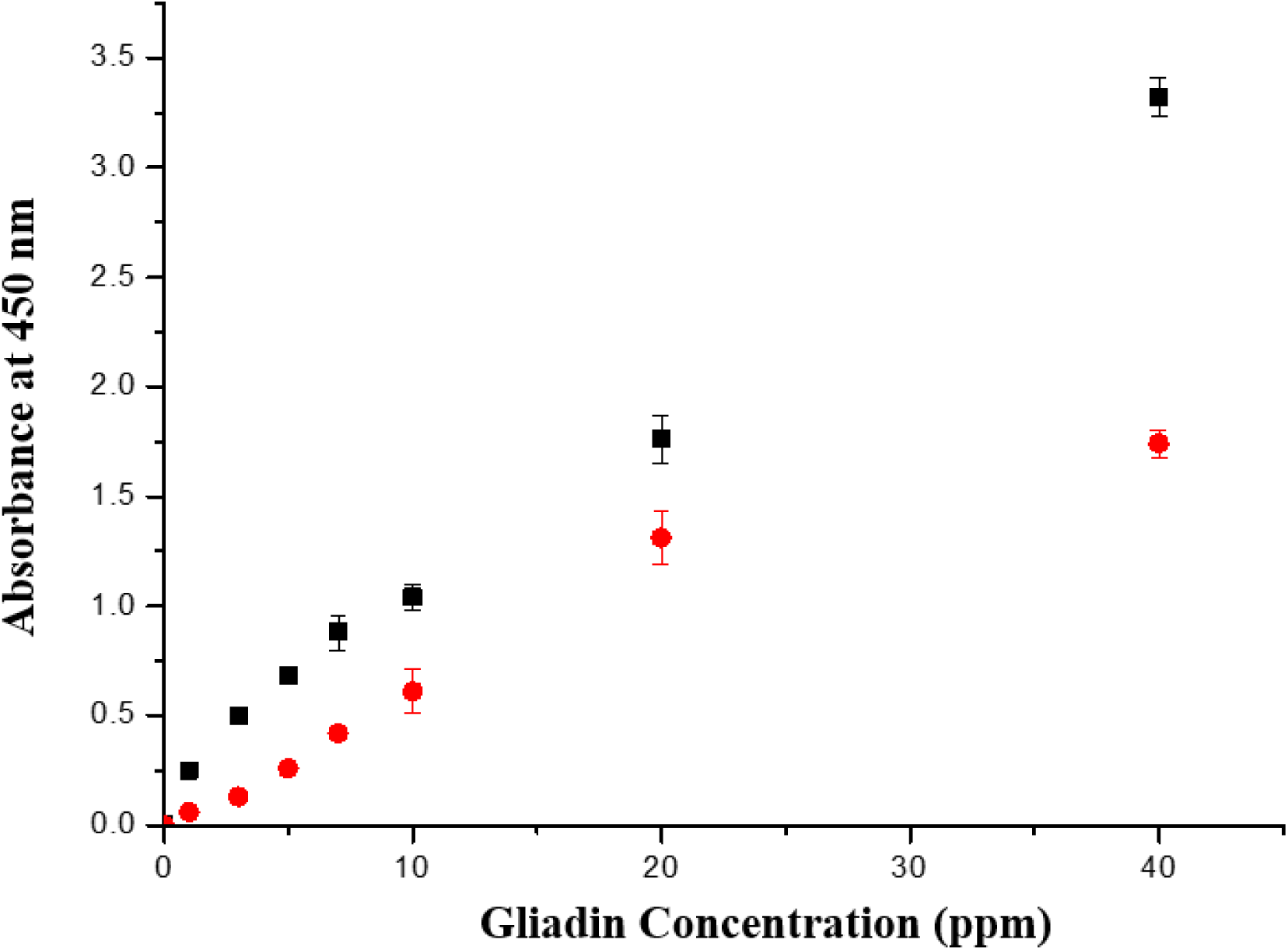
ELISA assay of gluten with 1% [C5DMIM][OMs] (black squares) and a 1% SDS solution (red dots).

**Figure 7.**
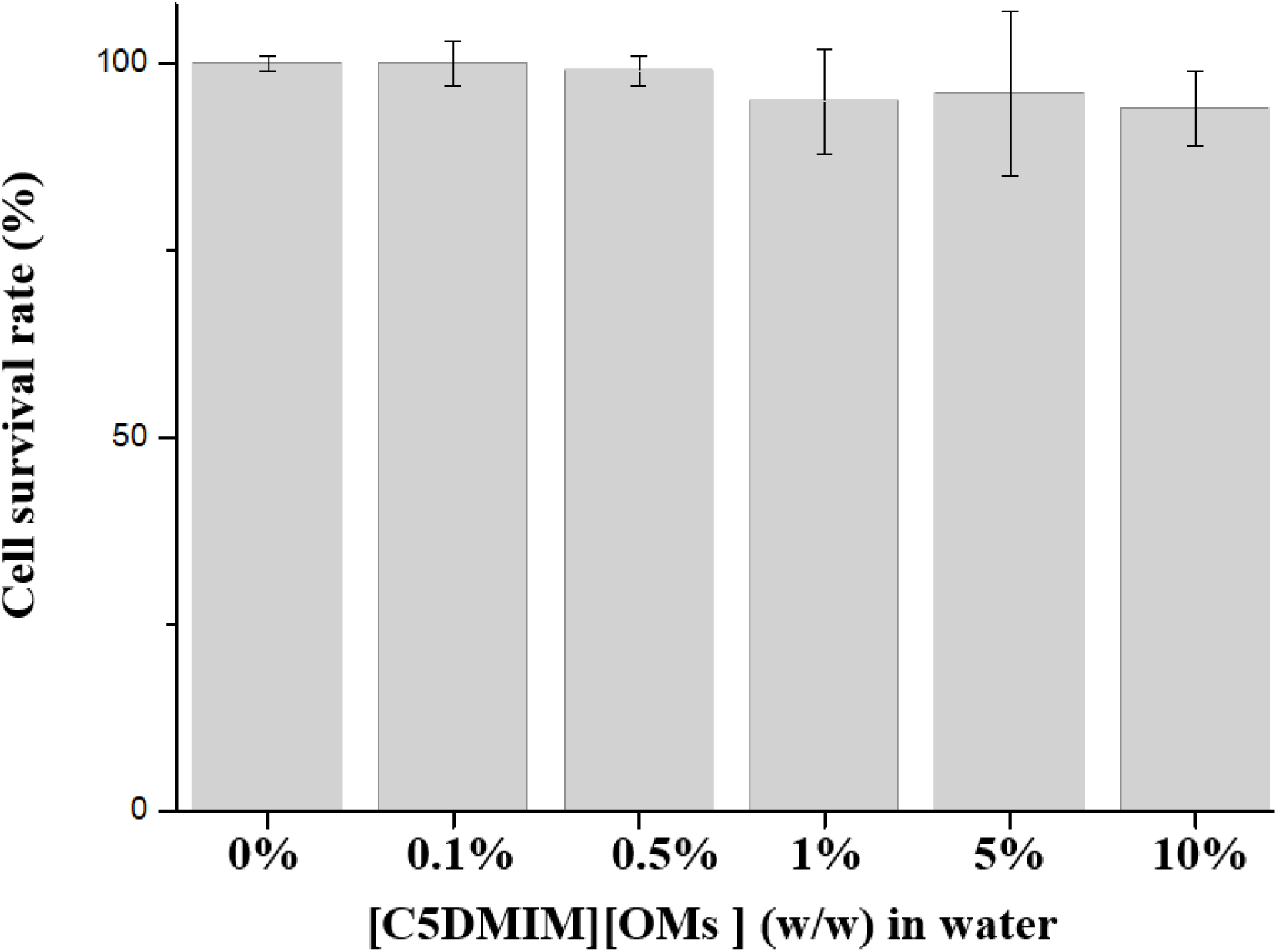
Cell toxicity with different concentrations of [C5DMIM][OMs]_aq_.

## Conclusions

Wheat is a major food for humans globally, but around 1%∼6% of people in the world have wheat allergy issues. Gluten from wheat can induce an inflammatory immune response in those patients. There are two major proteins of glutenin and gliadin in gluten. In recent years, some research claimed that gliadin is the major protein that induces the inflammatory immune response in patents. In food tests, gluten detection needs a lot of time for pretreating samples because gliadin is water insoluble. For this reason, we tried to synthesize an IL and applied it for rapid gluten extraction.

In this report, we synthesized ILs with different lengths of side chains and different anions of the imidazolium base IL to study the side chain and anion effects of imidazolium ILs applied to dissolve gliadin, and we purchased a gluten/gliadin ELISA kit to measure the extraction effect of imidazolium-based IL water solutions. As to the side chain effect of the gliadin solubility test, the ILs of [HDMIM], [TMIM], [C3DMIM], [C5DMIM], [C7DMIM], [C9DMIM], and [C12DMIM] with OM anions were synthesized to study gliadin solubility with an imidazolium base of a 1% IL water solution, and we found that the 1% IL water solution produced good gliadin solubility when the carbon chain was greater than 5 on imidazolium. In the anion effect test, we synthesized [C5DMIM] with different anions (F, Cl, NO_3_, HSO_4_, H_2_PO_4_, and OMs), and [C5DMIM] with OM anions produced the best gliadin solubility. In the kinetic curve of gliadin solubility, [C5DMIM][OMs] could dissolve more than 2500 ppm gliadin in 1 min and over 3000 ppm gliadin could be dissolved in the IL water solution in 3 min.

In this report, we measured the secondary structure of gliadin with and that without [C5DMIM][OMs] by CD spectroscopy, and found that the spectra did not significantly differ with or without [C5DMIM][OMs]. We measured the antibody-gliadin interaction by an ELISA after 1% [C5DMIM][OMs] extraction, and results showed good linearity of different concentrations of gliadin with [C5DMIM][OMs]. It showed that [C5DMIM][OMs] was not restricted by antibody binding with gliadin. In the recovery test, the average of recovery rate was 97.5% from rice noodles with gliadin added with 1% [C5DMIM][OMs].

Generally, the toxicity of cells will increase when the number of side chains increases. For this reason, we selected [C5DMIM][OMs] to test the cell toxicity, but we found no significant cell toxicity from [C5DMIM][OMs] in the N2a model. Those data showed that [C5DMIM][OMs] can be an extraction buffer for extracting gliadin from food. This technology can increase the effect of detecting gliadin and reduce organic solvents and multiple processes required to pretreat samples. It could be a great development to protect celiac patients.

## References

1. Costantino, A.; Aversano, G. M.; Lasagni, G.; Smania, V.; Doneda, L.; Vecchi, M.; Roncoroni, L.; Pastorello, E. A.; Elli, L. J. F. i. N., Diagnostic management of patients reporting symptoms after wheat ingestion. 2022, 9, 1007007.

2. Elli, L.; Branchi, F.; Tomba, C.; Villalta, D.; Norsa, L.; Ferretti, F.; Roncoroni, L.; Bardella, M. T. J. W. j. o. g. W., Diagnosis of gluten related disorders: Celiac disease, wheat allergy and non-celiac gluten sensitivity. 2015, 21 (23), 7110.

3. Gujral, N.; Freeman, H. J.; Thomson, A. B. J. W. j. o. g. W., Celiac disease: prevalence, diagnosis, pathogenesis and treatment. 2012, 18 (42), 6036.

4. Scherf, K. A. J. C. O. i. F. S., Immunoreactive cereal proteins in wheat allergy, non-celiac gluten/wheat sensitivity (NCGS) and celiac disease. 2019, 25, 35–41.

5. Serena, G.; D’Avino, P.; Fasano, A. J. F. i. N., Celiac disease and non-celiac wheat sensitivity: state of art of non-dietary therapies. 2020, 7, 152.

6. Siddiqui, K.; Uqaili, A. A.; Rafiq, M.; Bhutto, M. A. J. M., Human leukocyte antigen (HLA)-DQ2 and-DQ8 haplotypes in celiac, celiac with type 1 diabetic, and celiac suspected pediatric cases. 2021, 100 (11).

7. Vader, W.; Stepniak, D.; Kooy, Y.; Mearin, L.; Thompson, A.; van Rood, J. J.; Spaenij, L.; Koning, F. J. P. o. t. N. A. o. S., The HLA-DQ2 gene dose effect in celiac disease is directly related to the magnitude and breadth of gluten-specific T cell responses. 2003, 100 (21), 12390-12395.

8. Green, P. H.; Paski, S.; Ko, C. W.; Rubio-Tapia, A. J. G., AGA clinical practice update on management of refractory celiac disease: expert review. 2022.

9. Gromny, I.; Neubauer, K. J. I. J. o. E. R.; Health, P., Pancreatic Cancer in Celiac Disease Patients—A Systematic Review and Meta-Analysis. 2023, 20 (2), 1565.

10. Martín-Masot, R.; Herrador-López, M.; Navas-López, V. M.; Carmona, F. D.; Nestares, T.; Bossini-Castillo, L. J. I. J. o. M. S., Celiac Disease Is a Risk Factor for Mature T and NK Cell Lymphoma: A Mendelian Randomization Study. 2023, 24 (8), 7216.

11. Howdle, P. D. J. E. j. o. g.; hepatology, Gliadin, glutenin or both? The search for the Holy Grail in coeliac disease. 2006, 18 (7), 703–706.

12. de la Barca, A. M. C.; Yepiz-Plascencia, G. M.; Bφg-Hansen, T. C. J. L. s., Hydrophobic interactions between gliadin and proteins and celiac disease. 1996, 59 (23), 1951–1960.

13. Dziuba, M.; Dziuba, J.; Iwaniak, A. J. P. j. o. f.; sciences, n., Bioinformatics-aided characteristics of the structural motifs of selected potentially celiac-toxic proteins of cereals and leguminous plants. 2007, 57 (4), 405–414.

14. Chaudhary, N.; Virdi, A. S.; Dangi, P.; Khatkar, B. S.; Mohanty, A. K.; Singh, N. J. F. H., Protein, thermal and functional properties of α-, γ-and ω-gliadins of wheat and their effect on bread making characteristics. 2022, 124, 107212.

15. Solé-Jamault, V.; Davy, J.; Cochereau, R.; Boire, A.; Larré, C.; Denery- Papini, S. J. J. o. C.S., Optimization of large-scale purification of omega gliadins and other wheat gliadins. 2022, 103, 103386.

16. Watanabe, C.; Ando, Y.; Kumagai, H.; Miyamoto, M.; Fujita, Y.; Kato, M.; Nakayama, M.; Maruyama, N.; Yamagata, T.; Yoshihara, S. J. I. J. o. P., A Pediatric Case of Wheat-Dependent, Exercise-Induced Anaphylaxis Solely Associated with High-Molecular-Weight Glutenin. 2022, 89 (9), 937–937.

17. Zhang, Y.; Zhang, X.; Zhang, Z.; Chen, Z.; Jing, X.; Wang, X. J. L., Effect of high hydrostatic pressure treatment on the structure and physicochemical properties of millet gliadin. 2022, 154, 112755.

18. Urade, R.; Sato, N.; Sugiyama, M. J. B. r., Gliadins from wheat grain: An overview, from primary structure to nanostructures of aggregates. 2018, 10, 435–443.

19. Di Cagno, R.; De Angelis, M.; Lavermicocca, P.; De Vincenzi, M.; Giovannini, C.; Faccia, M.; Gobbetti, M. J. A.; microbiology, e., Proteolysis by sourdough lactic acid bacteria: effects on wheat flour protein fractions and gliadin peptides involved in human cereal intolerance. 2002, 68 (2), 623–633.

20. Gerez, C. L.; Dallagnol, A.; Rollán, G.; de Valdez, G. F. J. F. m., A combination of two lactic acid bacteria improves the hydrolysis of gliadin during wheat dough fermentation. 2012, 32 (2), 427–430.

21. Wagh, S. K.; Gadge, P. P.; Padul, M. V. J. P.; proteins, a., Significant hydrolysis of wheat gliadin by Bacillus tequilensis (10bT/HQ223107): a pilot study. 2018, 10, 662-667.

22. Wieser, H. J. A. P., Relation between gliadin structure and coeliac toxicity. 1996, 85, 3-9.

23. Gómez Castro, M. F.; Miculán, E.; Herrera, M. G.; Ruera, C.; Perez, F.; Prieto, E. D.; Barrera, E.; Pantano, S.; Carasi, P.; Chirdo, F. G. J. F. i. i., P31–43 gliadin peptide forms oligomers and induces NLRP3 inflammasome/caspase 1-dependent mucosal damage in small intestine. 2019, 10, 31.

24. Tague, E. J. J. o. t. A. C. S., The iso-electric points of gliadin and glutenin. 1925, 47 (2), 418–422.

25. Arnold, L. K.; Choudhury, R.; Dangoria, D. C. In The solubility of wheat gluten in various aqueous solutions, Proceedings of the Iowa Academy of Science, 1964; pp 193–196.

26. Fu, B.; Sapirstein, H.; Bushuk, W. J. J. o. C. S., Salt-induced disaggregation solubilization of gliadin and glutenin proteins in water. 1996, 24 (3), 241–246.

27. Popineau, Y.; Godon, B., Surface hydrophobicity of gliadin components. 1982.

28. Eyckens, D. J.; Henderson, L. C. J. F. i. c., A review of solvate ionic liquids: Physical parameters and synthetic applications. 2019, 7, 263.

29. Lei, Z.; Chen, B.; Koo, Y.-M.; MacFarlane, D. R. J. C. R., Introduction: ionic liquids. ACS Publications: 2017; Vol. 117, pp 6633–6635.

30. Vander Hoogerstraete, T.; Onghena, B.; Binnemans, K. J. T. j. o. p. c. l., Homogeneous liquid–liquid extraction of metal ions with a functionalized ionic liquid. 2013, 4 (10), 1659–1663.

31. Ventura, S. P.; e Silva, F. A.; Quental, M. V.; Mondal, D.; Freire, M. G.; Coutinho, J. A. J. C. R., Ionic-liquid-mediated extraction and separation processes for bioactive compounds: past, present, and future trends. 2017, 117 (10), 6984–7052.

32. Patel, R.; Kumari, M.; Khan, A. B. J. A. b.; biotechnology, Recent advances in the applications of ionic liquids in protein stability and activity: a review. 2014, 172, 3701–3720.

33. Yang, W.; Dunlap, J. R.; Andrews, R. B.; Wetzel, R. J. H. m. g., Aggregated polyglutamine peptides delivered to nuclei are toxic to mammalian cells. 2002, 11 (23), 2905–2917.

34. Yushchenko, T.; Deuerling, E.; Hauser, K. J. B. J., Insights into the aggregation mechanism of PolyQ proteins with different glutamine repeat lengths. 2018, 114 (8), 1847–1857.

35. Mendonça, C. M.; Balogh, D. T.; Barbosa, S. C.; Sintra, T. E.; Ventura, S. P.; Martins, L. F.; Morgado, P.; Filipe, E. J.; Coutinho, J. A.; Oliveira, O. N. J. P. C. C. P., Understanding the interactions of imidazolium-based ionic liquids with cell membrane models. 2018, 20 (47), 29764–29777.

36. Yoo, B.; Jing, B.; Jones, S. E.; Lamberti, G. A.; Zhu, Y.; Shah, J. K.; Maginn, E. J. J. S. r., Molecular mechanisms of ionic liquid cytotoxicity probed by an integrated experimental and computational approach. 2016, 6 (1), 19889.

37. Montalbán, M.; Bolívar, C.; Diaz Banos, F. G.; Víllora, G. J. J. o. C.; Data, E., Effect of temperature, anion, and alkyl chain length on the density and refractive index of 1-alkyl-3-methylimidazolium-based ionic liquids. 2015, 60 (7), 1986-1996.

